# Molecular signature of domestication in the arboviral vector *Aedes aegypti*

**DOI:** 10.1101/2023.03.13.532092

**Authors:** A.N. Lozada-Chávez, I. Lozada-Chávez, N. Alfano, U. Palatini, D. Sogliani, S. Elfekih, T. Degefa, M.V. Sharakhova, A. Badolo, S. Patchara, M. Casas-Martinez, B.C Carlos, R. Carballar-Lejarazú, L. Lambrechts, J.A. Souza-Neto, M. Bonizzoni

**Author notes:** N.A.: Human Technopole, Viale Rita Levi-Montalcini 1, 20157, Milan, Italy U.P: Laboratory of Neurogenetics and Behavior, The Rockefeller University, New York, NY, USA RCL: Department of Microbiology & Molecular Genetics, University of California, Irvine, CA 92697-4025, USA B.C.C.: São Paulo State University (UNESP), School of Agronomy, Crop Protection Department, Research group on Integrated pest management (AGRIMIP), Botucatu, Brazil; J.A.S.N.: Department of Diagnostic Medicine/Pathobiology, College of Veterinary Medicine, Kansas State University, Manhattan, Kansas, USA.

## Abstract

**Background:** Domestication is a complex, multi-stage and species-specific process that results in organisms living close to humans. In the arboviral vector *Aedes aegypti* adaptation to living in proximity with anthropogenic environments has been recognized as a major evolutionary shift, separating a generalist form, *Aedes aegypti formosus* (Aaf), from the domestic form *Aedes aegypti aegypti* (Aaa), which tends to deposit eggs artificial containers and bite humans for a blood meal. These behaviors enhance the mosquito vectorial capacity. The extent to which domestication has impacted the *Ae. aegypti* genome has not been thoroughly investigated yet.

**Results:** Taking advantage of two forms’ distinct and historically documented geographic distributions, we analyzed the genomes of 634 worldwide *Ae. aegypti* mosquitoes. Using more than 300 million high-confidence SNPs, we found a unique origin for all out-of-Africa *Ae. aegypti* mosquitoes, with no evidence of admixture events in Africa, apart from Kenya. A group of genes were under positive selection only in out-of-Africa mosquitoes and 236 genes had nonsynonymous mutations, occurring at statistically different frequencies in Aaa and Aaf mosquitoes.

**Conclusion:** We identified a clear signal of genetic differentiation between Aaa and Aaf, circumscribed to a catalogue of candidate genes. These “*Aaa molecular signature*” genes extend beyond chemosensory genes to genes linked to neuronal and hormonal functions. This suggests that the behavioral shift to domestication may rely on the fine regulation of metabolic and neuronal functions, more than the role of a few significant genes. Our results also provide the foundation to investigate new targets for the control of *Ae. aegypti* populations.

## Background

The complex and multistage process that brings animals to live in proximity with anthropogenic environments has had a tremendous impact on both animals and human evolution since the Neolithic time when humans started to breed animals as food or commodity sources, protection, transportation, and company (1). For animals such as sheep, goats, cattle, chinchilla, American minks and shrimp domestication has been a human-driven process (2). For other species, domestication has been a self-selective natural process, which has resulted in an inherited attraction and adaptation to anthropogenic environments (2). A recognized consequence of domestication is exposure to zoonotic diseases because the domesticated animal may act as a reservoir or an amplifier of pathogens mainly acquired through interaction with wildlife (3).

The primary vector of arboviruses worldwide, the mosquito *Aedes aegypti*, exists as two different subspecies: the generalist *Aedes aegypti formosus* (Aaf) and the domesticated *Aedes aegypti aegypti* (Aaa). The distinction between the two subspecies has epidemiological relevance because Aaa tends to have a higher vectorial capacity than Aaf mosquitoes due to its behavior patterns of domestication, such as aptitude to oviposit in clean water of artificial containers and preference to blood feed on humans, and its higher vector competence for arboviruses (4–6). The two subspecies are distinguished at the phenotypic level, with Aaf a darker body color than Aaa (7,8). However, body color is not a binary phenotype, resulting in uncertainty (8,9).

*Aedes aegypti* is native of Africa and diverged from its closest relative, *Aedes mascarensis,* between 4 to 15 million years ago (MYA) (10). Nowadays, *Ae. aegypti* can be found throughout the tropical and subtropical regions of the world, but its geographic populations are not homogenous in terms of domesticated behaviors. Out-of-Africa populations, which originated through the transatlantic slave trade from African populations, preferentially bite humans and lay eggs in freshwater of human-made containers (11). African populations tend to be generalist. While it is debatable when and which are the ecological drivers that caused this shift (6,12,13), genetic data based on microsatellite markers and exome sequencing have revealed a clear genetic differentiation between the two morphological subspecies, supporting a single sub-speciation event that probably occurred in West Africa, with the absence of gene flow between out-of-Africa domesticated and African generalist mosquitoes for at least 500 years (6,10,13–17). However, in a few African locations such as Kenya, Angola and urban sites of West Africa, *Ae. aegypti* mosquitoes that preferentially bite humans are sampled, suggesting recent reintroductions from out-of-Africa mosquitoes leading to admixture, persistence of descendants from the ancestral Aaa population of West Africa or incipient and independent domestication events (6,12,16,18).

It has long been speculated that domestication has strong genomic bases in *Ae. aegypti* because this species appears to have a high genetic diversity on a micro-geographic scale, and it is known to be fast evolving (16,19–21). However, the extent of the molecular differentiation in the genomes of the two subspecies in relation to biological functions associated with the behavioral switch to domestication have not been extensively investigated yet. Most efforts have been focused on identifying specific genes linked to hallmarks of domestication such as host-seeking behavior or insecticide resistance, the latter being a side effect of recurrent insecticidal interventions to control mosquito populations (22–24), based on the differential expression across *Ae. aegypti* populations (25–28) and/or the presence of nonsynonymous variants within a few target loci, including genes encoding for detoxification enzymes (*e.g.*, P450s) (23), neurotransmitter receptors (*e.g.*, Ir8) (29), the para sodium channel gene (*e.g.*, *kdr* mutations) (23,30,31), and odorant receptors (*e.g.*, AeOR4) (6).

Here, we present a comprehensive screening of genomic features in *Ae. aegypti* associated with adaptations to microgeographic and expected/recorded domesticated behaviors after its subspeciation event from the comparative analysis of the genomes of 123 out-of-Africa and 511 African mosquitoes. Among these genomes, we identified more than 300 million high-confidence SNPs, which we used to assess population structure and test for signatures of molecular evolution. Our analyses led to circumscribing the genetic basis of domestication to a catalogue of candidates, which are strongly differentiated between Aaa and Aaf mosquitoes.

## Results

### 300 million high confidence SNPs detected across 634 worldwide *Ae. aegypti* genomes

We analyzed the complete genome of 694 *Aedes* spp. mosquitoes, including 4 *Ae. mascarensis* and 21 *Ae. albopictus* samples. After removing datasets with low-quality mapping or molecular identification as either *Ae. albopictus* or *Aedes simpsoni*, we obtained a final dataset of 634 *Ae. aegypti* genomes divided into two major geographical groups: African samples and out-of-Africa samples (**Table 1, Fig. 1)**. Based on the presence or absence of the *Nix* gene, 192 and 395 mosquitoes were classified as males and females, respectively. Since the DNA of sperms can be stored in female spermatheca, the remaining 47 individuals with partial coverage (2-44%) of the *Nix* gene were considered females.

**Fig. 1.**
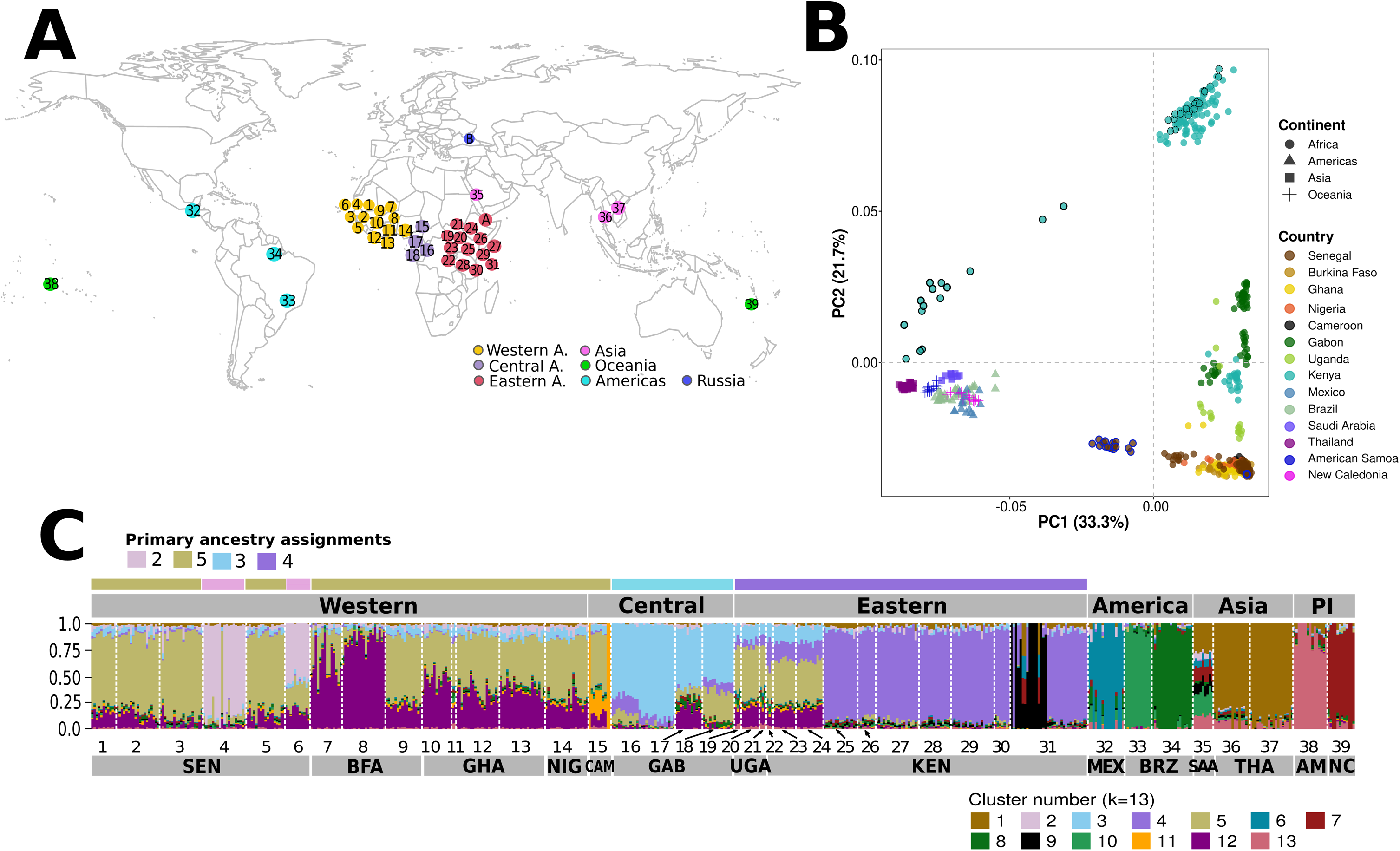
Worldwide population structure of African and out-of-Africa samples of *Aedes aegypti*. (**A**) Map of worldwide collection sites of *Aedes aegypti* populations used in this study. Sites are color-coded by region, indicating the sampled regions and the number of their position according to Table 1. (**B**) A Principal Component Analyses (PCA) generated with a panel of 1.5 million of non-redundant and biallelic SNPs from the initial dataset of 634 samples. Each dot represents an individual being color-coded by country (filled circles) and continent (different symbols). Samples from RAB (Kenya) are distinguished as RABd (black solid outlined circles) and RABs (black dotted outlined circles), whereas samples from NGY (Senegal) are divided by showing a strong affinity to Out-of-Africa (blue solid outlined circles) or Western African populations (blue dotted outlined circles). (**C**) ADMIXTURE analyses of population structure for all sampled populations with k=13. On the Y-axis, each vertical bar represents the probability (Q-values from 0 to 1) of assignment of a single individual to each genetic cluster, and each population is separated by a vertical white line and colored according to the legend at the bottom. Individuals with mixed ancestry are denoted by bars with different colors. On the X-axis, country names and numbers are reported according to the abbreviation described in Table 1. African populations were ordered based on their geographical location (Western, Central, and Eastern), and Out of Africa by their corresponding continent. Based on their primary ancestry assignments, African populations are grouped in four genetic clusters: Western (THI and NGY, cluster 2), Western-Central (cluster 5), Central (cluster 3), and Eastern (cluster 4) Africa.

**Table 1.**
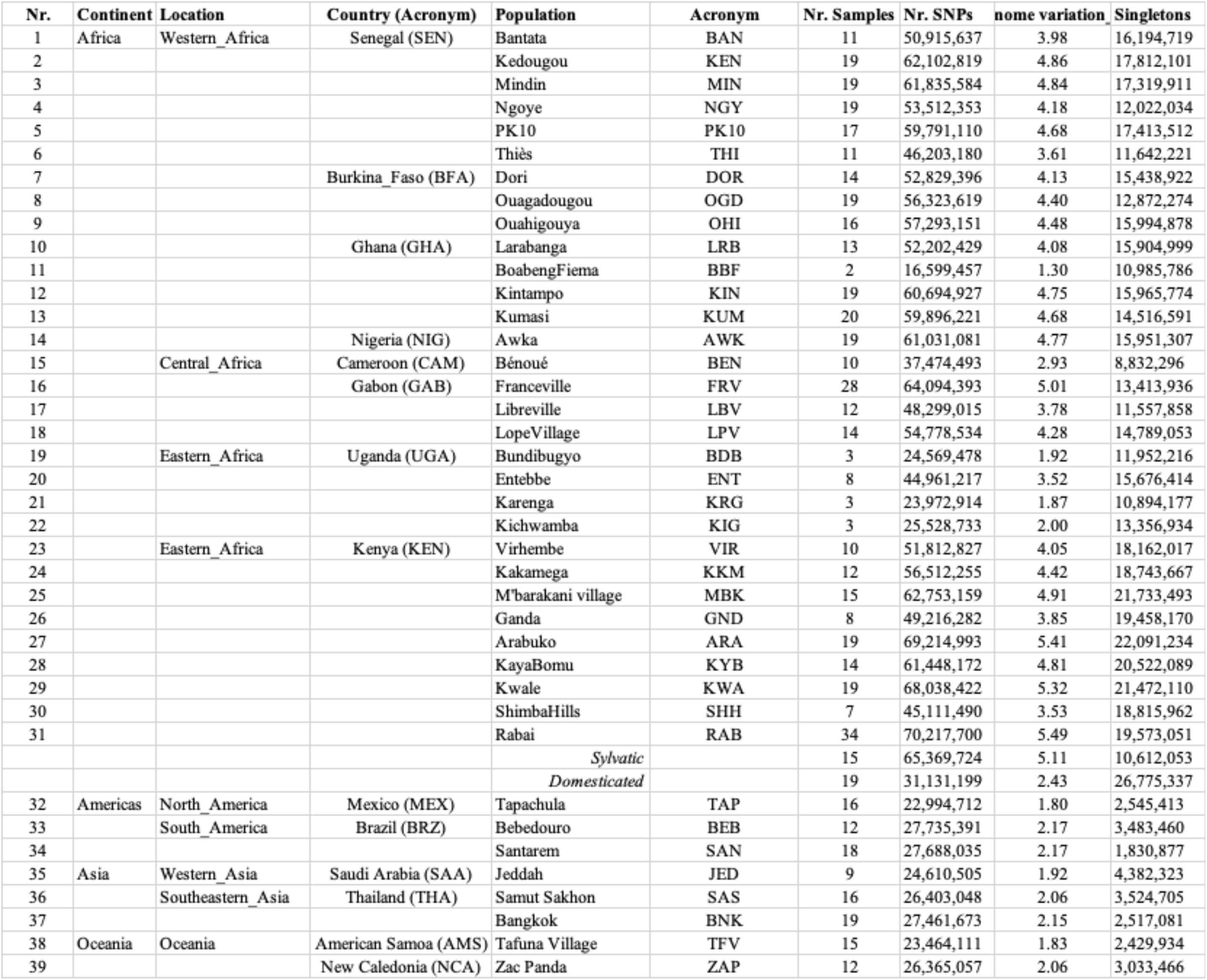
(should be added to page 5, line 121) List of the Aedes aegypti populations analyzed, including the number of sequenced genomes (Nr. samples), the total number of SNPs identified (Nr. SNPs), the percentage of variation across the genome and the number of singletons.

We identified a total of 322,552,899 high-confidence SNPs across the 634 *Ae. aegypti* genomes by using two variant callers to recalibrate of our initial predicted SNPs dataset (∼207 million) and ∼485 thousand SNPs from ten published datasets (see Methods). We found no statistical difference in the number of SNPs between males and females (t-test=2.75, p>0.05), while SNPs number correlated with the length of chromosomes (length of chromosome 2: 2473 Mbs, 44.88% of total number of SNPs; length of chromosome 3: 409 Mbs, 37.71% of total number of SNPs and length of chromosome 1: 310 Mbs, 26.40 % of total number of SNPs, rank coefficient correlation > 0.87, p<0.005). SNP distribution does not fit a model of “random distribution” across the chromosomes (chi-square test, p>0.1), but SNP density is highest at the telomere and decreases towards the centromere in all three chromosomes. Due to the highly repetitive nature of the *Ae. aegypti* genome (32), we distinguished between SNPs located in repetitive (R-SNPs) and non-repetitive (NR-SNPs) sequences. As expected, the number of biallelic and multiallelic R-SNPs (223 million) is higher in comparison to NR-SNPs (91 million), with no significant differences found in these trends for African and out-of-Africa populations (t-test=1.3, p>0.05). NR-SNPs are mostly located in intergenic regions (48%) and introns (43%), and to a lesser extent within exons (9%); whereas R-SNPs are equally distributed across introns and intergenic regions. Further analyses in this study are focused on the NR-SNPs dataset.

Among tested genomes, we identified 95 close relatives that represent >50% of individuals from six African populations, including Thiès (THI), Bundibugyo (BDB), Kichwamba (KIC), Virhembe (VIR), Bénoué (BEN), and Rabai (RAB), leaving a dataset of 539 *Ae. aegypti* genomes for which we performed a Principal Component Analysis (PCA). We recovered four well defined clusters representing three metapopulations from Africa (Western, Central-Eastern, and Eastern regions), and one from non-African populations, but we did not identify any cluster with mosquitoes that were previously described as “domesticated” in Rabai (6). A second PCA including all 634 mosquitoes showed RAB-related individuals are those with a closer affiliation to out-of-Africa mosquitoes (**Fig. 1B**). These results highlight that “domesticated” Rabai mosquitoes are inbred. Based on these results, we included all individuals from RAB, subdividing them into the “domesticated” (RABd) or “feral” (RABs) group, resulting in a final dataset of 554 *Ae. aegypti* genomes.

### High genetic diversity and rare variants are more common in African populations

Among the 554 *Ae. aegypti* genomes, we detected 314,365,358 high-confidence SNPs; 81% are biallelic and 19% multiallelic. The average number of variants (R-SNPs, NR-SNPs) *per* population (46±16 million) represents 3.60% of the total variation detected in the genome, with a difference between African (3.99%) and out-of-Africa (2.02%) populations (**Table 1**). This can be explained by the presence of almost two-fold more variants in African (51.91±13.72 million) than in out-of-Africa (25.84±1.90 million) populations (unpaired Wilcoxon signed rank test, p=0.000196), as previously observed (13, 18). Within African populations, the Western (SEN, BFA, GHA) and Eastern (KEN) regions have a similar number of variants (56.51±11.69 and 56.06±11.94 million, respectively), which is higher than that detected in Central populations (51.16±11.19 million). Conversely, out-of-Africa populations not only have less variants (25.84±1.90 million), but also less variance in the number of variants across them.

Mean nucleotide diversity (*π*) and Tajima’s D (*D*) estimates were calculated with a non-overlapping sliding window of 10kb across chromosomes and populations to identify demographic changes. On average, *π* is found to be higher in African (0.67±0.33) than out-of-Africa (0.34±0.09) populations. Also, populations from Central and West Africa have more genome intervals with negative Tajima’s D values on each chromosome, which are more concentrated towards the telomeres (63% of all sliding windows). Both estimates indicate that high genetic diversity and rare variants are more common across African populations, suggesting novel mutations due to population expansions. Conversely, NGY, RABd and out-of-Africa populations have more genome intervals with positive Tajima’s D values, along with a reduced number of rare variants and nucleotide diversity (lower π values), suggesting a pervasive effect of bottlenecks or inbreeding due to one or repeated population contractions. These results and trends are robust to estimates with different sliding window sizes.

### Unique origin for out-of-Africa mosquitoes and no evidence of incipient domestication or current admixture events between African and out-of-Africa populations beyond Kenya

Using a set of 1,530,512 biallelic SNPs that are shared across 80% of the 554 genomes, with no linkage disequilibrium and a minor allele frequency (MAF) of 0.01, we performed admixture analyses (K=3-13) (**Fig. 1C**) and compared pairwise fixation indexes (Fst) as a measurement of population genetic divergence and evaluating the presence of local introgression. Additionally, we identified phylogenetic relationships among ‘individuals’ and ‘populations’ with two independent maximum likelihood (ML) trees that were reconstructed using exomic biallelic SNPs and *Ae. albopictus* as an outgroup (**Fig. 2**). Admixture analysis identified five well-defined clusters, representing one group from out-of-Africa populations and four African metapopulations from the Western (clusters 2), Western/Central (cluster 5), Central (cluster 3) and Eastern (cluster 4) regions (**Fig. 1C**). Furthermore, genetic separation is observed between populations west (Uganda, Western Kenya) and east (Eastern Kenya) of the Rift Valley. The human-feeding mosquitoes of the Senegalese NGY and THI populations (within cluster 2) and RABd (within cluster 9) formed distinct clusters. In the ML-phylogeny of individuals, RABd was found to form a single well-supported branch with JED mosquitoes (bootstrap=70%), as previously observed (10). This finding is supported by the lowest genetic divergence found between RABd and JED (Fst__RABd-JED_=0.147±0.01) in comparison to the divergence found against other out-of-Africa (Fst__RABd-Aaa_=0.179±0.091, range=0.156-0.214) and African populations (Fst__RABd-Aaf_=0.183±0.013, range=0.148-0.231).

**Fig. 2.**
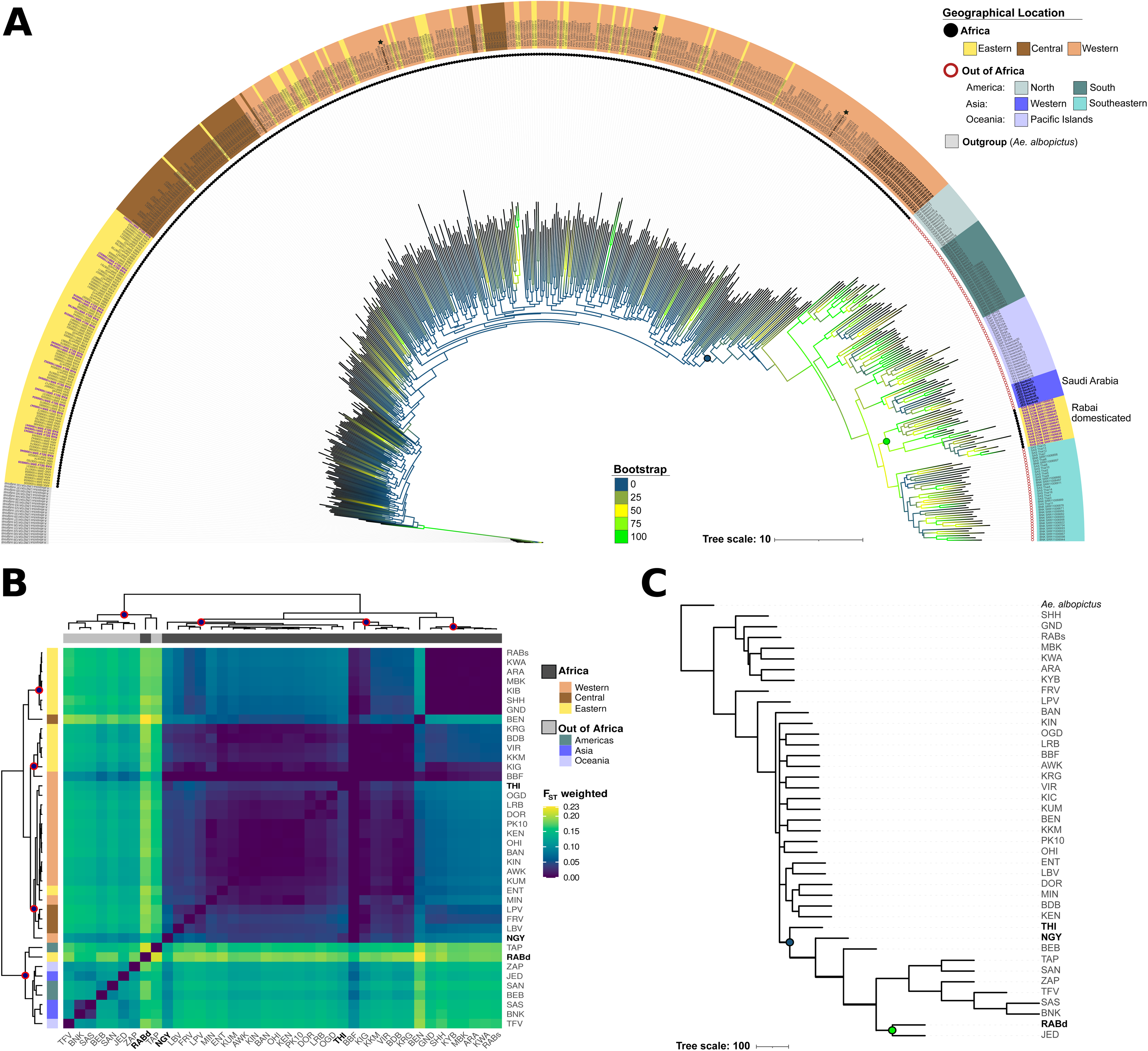
Evolutionary relationships among 554 *Ae. aegypti* genomes from 39 worldwide populations. (**A**) A maximum likelihood (ML) tree for individuals including 554 mosquitoes, after 1,000 bootstraps resampling of the dataset. RABd is colored in purple and bold font. THI and NGY mosquitoes are colored in black and bold font. Bootstraps support for each relationship, as well as the geographical region of sampling for each individual, are color-coded accordingly to their symbology. Filled black circles at the base of each individual name are denoting Africa individuals, while open dark circles represent out-of-Africa mosquitoes. **(B)** Divergence among populations based on weighted Fst-based genetic distances, according to the Weir and Cockerham approach. The heatmap is showing a *complete* hierarchical clustering of Fst values from pairwise comparisons across all populations, by using a *euclidean* distance and 1,000 bootstrap replicates. On both axes, population names are shown according to the abbreviation listed on Table 1, with populations clustered and colored by country (Y-axis) and geographical region (X-axis). The diagonal in the matrix represents the comparison with the same population (zero difference, in black), while the degree of divergence for each comparison is shown according to the color symbology at the right bottom. (**C**) A ML tree for populations reconstructed using the same SNPs dataset of tree of individuals but estimating SNPs allele frequencies after 1,000 bootstraps resampling of the dataset. *Ae. albopictus* was used in both ML-trees as an outgroup, and branch lengths in both ML-tress are proportional to the amount of genetic divergence that has occurred.

The ML-phylogeny for individuals (**Fig. 2A**) also identified a single well-defined branch including THI and NGY mosquitoes with all out-of-Africa populations (bootstrap <50%). The branch length of the nodes grouping THI and NGY is larger (7.27±0.66) than that grouping non-African mosquitoes altogether (4.45±1.24). Both THI and NGY show higher divergence with out-of-Africa populations (Fst__THI-Aaa_: 0.127±0.12, range 0.113-0.175; Fst__NGY-Aaa_: 0.104±0.12, range: 0.128-0.16), in comparison to the divergence found against other African populations (Fst__THI-Aaf_ : 0.048±0.025, range 0.004–0.02 and Fst__NGY-Aaf_: 0.055±0.021, range 0-0.081). Additionally, Fst values estimated for THI and NGY with respect to mosquitoes from America, Asia and Oceania, which were colonized in a subsequent manner (11), showed a consistent high variance, whereas within genetic divergence in both THI and NHY populations was low (NGY: F_IS_=0.015, THI: F_IS_=0.013), with only one THI and two NGY mosquitoes grouping with Central Africa populations. Apart from NGY and THI, we observed grouping of mosquitoes within Africa in sampled populations from West and East Africa, with extensive intermixing (*i.e.*, interleaving branches) in Central Africa, and no distinct separation among mosquitoes sampled in urban or rural/forestry sites, suggesting frequent gene flow in mosquitoes from these habitats. Results obtained from the ML-phylogeny for individuals were further confirmed by both the ML-populations phylogeny based on allele frequencies (33) and the F3 statistic, which tested genetic admixture among African populations and between African and out-of-Africa populations (34). THI, NGY and all out-of-Africa populations were found to form a single well-supported branch in the ML-populations phylogeny (bootstrap= 78%) (**Fig. 2C**). The phylogenetic relationship of both THI and NGY with out-of-Africa populations is not the product of admixture events, given that the F3 test was rejected in all cases (Z-cores values > −3.0). The same result was obtained when extending the F3 statistics to other African populations. On the contrary, there is evidence of admixture among African populations from geographically close regions, independently of their urban or forest location. The ML-populations phylogeny also showed that the amount of genetic drift is higher across out-of-African populations, in agreement with our findings of low genetic diversity and positive values of Tajima’s D.

### Signatures of local genetic adaptation is detected in out-of-Africa mosquitoes

Admixture and phylogenetic analyses confirmed the genetic separation between African mosquitoes and domesticated out-of-Africa samples and highlighted RABd, NGY and THI as the African samples most closely related to Aaa mosquitoes. To further identify patterns of molecular differentiation among our samples, we screened for outlier SNPs using the program *PCAdapt* (35). Among the 10,030 outliers detected, the majority were located within intronic (45%) and intergenic (26%) regions, while 27% mapped to protein-coding exons of 2,266 genes. In agreement with results from our phylogenetic analyses, apart from THI, NGY and RABd, all African populations grouped together and were separated from out-of-Africa mosquitoes (**Fig. 3**). The first three principal components (PCs) accounted for 95% of the data (one sample t-test for PCs, *p*<0.001; pairwise t-test for populations, *p*<0.001). PC1 captured 6,041 outliers distributed across 1639 genes that support local adaptations for THI, NGY, RABd, RABs and out-of-Africa populations; PC2 and PC3 included 3,559 outliers in 986 genes highlighting local adaptations in NGY and eight Kenyan populations. The remaining PCs explain 5% of the data and identified 429 outliers across 233 genes indicating local adaptations in out-of-Africa populations.

**Fig. 3.**
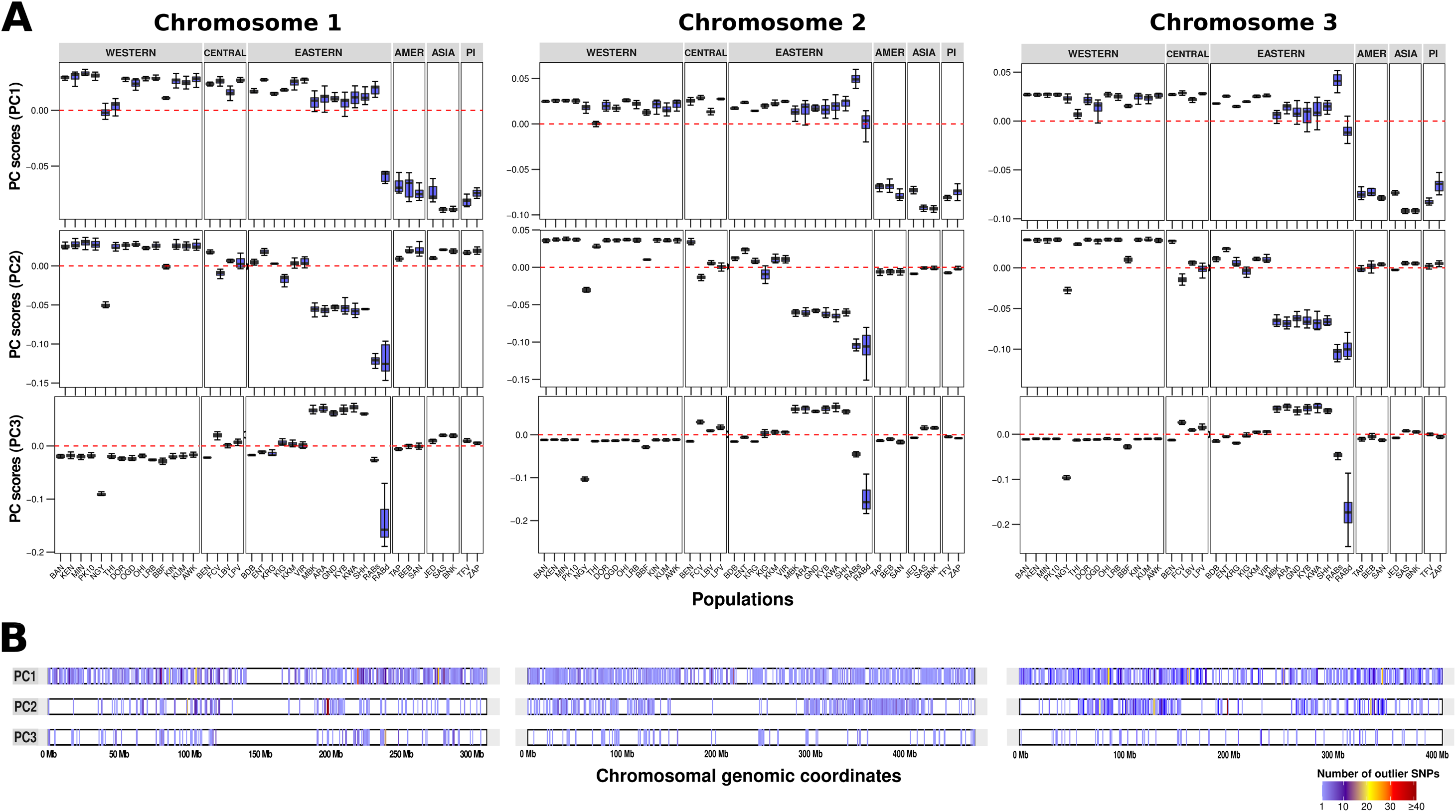
Genomic signatures of local adaptation. (**A**) Distribution of the PC scores estimated from outlier SNPs across the three chromosomes of *Ae. aegypti*. On each chromosome panel, the PC scores for each population (on the X-axis) are shown for PC1, PC2, and PC3 (on the Y-axis). Each PC highlights different genomic variants associated to local adaptation in Aaa populations relative to their Aaf counterparts. Each boxplot depicts the variation of the data in a population, indicating the first quantile, mean, third quantile, and the lower and upper whiskers as minimum and maximum variation of the data population, respectively. PC scores equal to zero are denoted with a horizontal dotted red line. Population names are reported according to their abbreviation in Table 1. (**B**) Distribution of 2,266 protein-coding genes across the *Ae. aegypti* genome that are harboring at least one of the genomic variants detected in (A). The 2,266 genes estimated as locally adapted were plotted for each PC and panel chromosome (1 in left; 2 in central; 3 in right). Chromosomes are depicted as horizontal bars, with locally adapted genes denoted as vertical lines, the frequency of outlier SNPs that each gene harbor is represented by a color gradient according to the symbology. Approximate genomic positions for each chromosome are shown at the bottom.

The hallmark of domestication in *Ae. aegypti* is higher attraction to humans than animals, a phenotype that was used to classify Aaa and Aaf mosquitoes, prior to the discovery of Aaf mosquitoes preferentially blood-feeding on humans in Cape Verde and Senegal, and is regulated by chemosensory receptors (6,26,36). Among the 198 *Ae. aegypti* chemosensory genes, 32 had outliers. Relative to other functional categories, most genes with outliers were associated with functions such as protease activity (35 genes out of 292 genes), detoxification (29 genes out of 198 genes), immunity (62 out of 391 genes) or biosynthetic and metabolic processes, signaling and receptor activities, metal ion homeostasis and (GTPase) binding activities. All genes harboring outliers showed high genetic differentiation (sites average Fst≥0.09) and deviation from neutrality (Tajima’s D: one sample t-test, m=0, *p*<0.001), suggesting signals of selection. Additionally, outliers in 306 genes resulted in nonsynonymous mutations, which occurred at a significantly different allele frequency in African and out-of-African mosquitoes in 236 genes, including the odorant receptor (OR) genes *OR91* and *OR86*, the ionotropic receptors (Ir) *Ir41g*, *7g* and *8a*; the gustatory receptor (GR) *9*; immunity genes as *Toll5A,* the gram-negative binding protein (GNBP) A2, the clip-domain serine proteases *CLIPE12*, and *CLIPE8*, and several genes with unknown functions (*i.e*. AAEL025393, AAEL001559, AAEL020878, AAEL022804, AAEL022666, AAEL019451, AAEL012825, AAEL012783, AAEL012268, AAEL010998, AAEL008698) (**Fig. 4**).

**Fig. 4.**
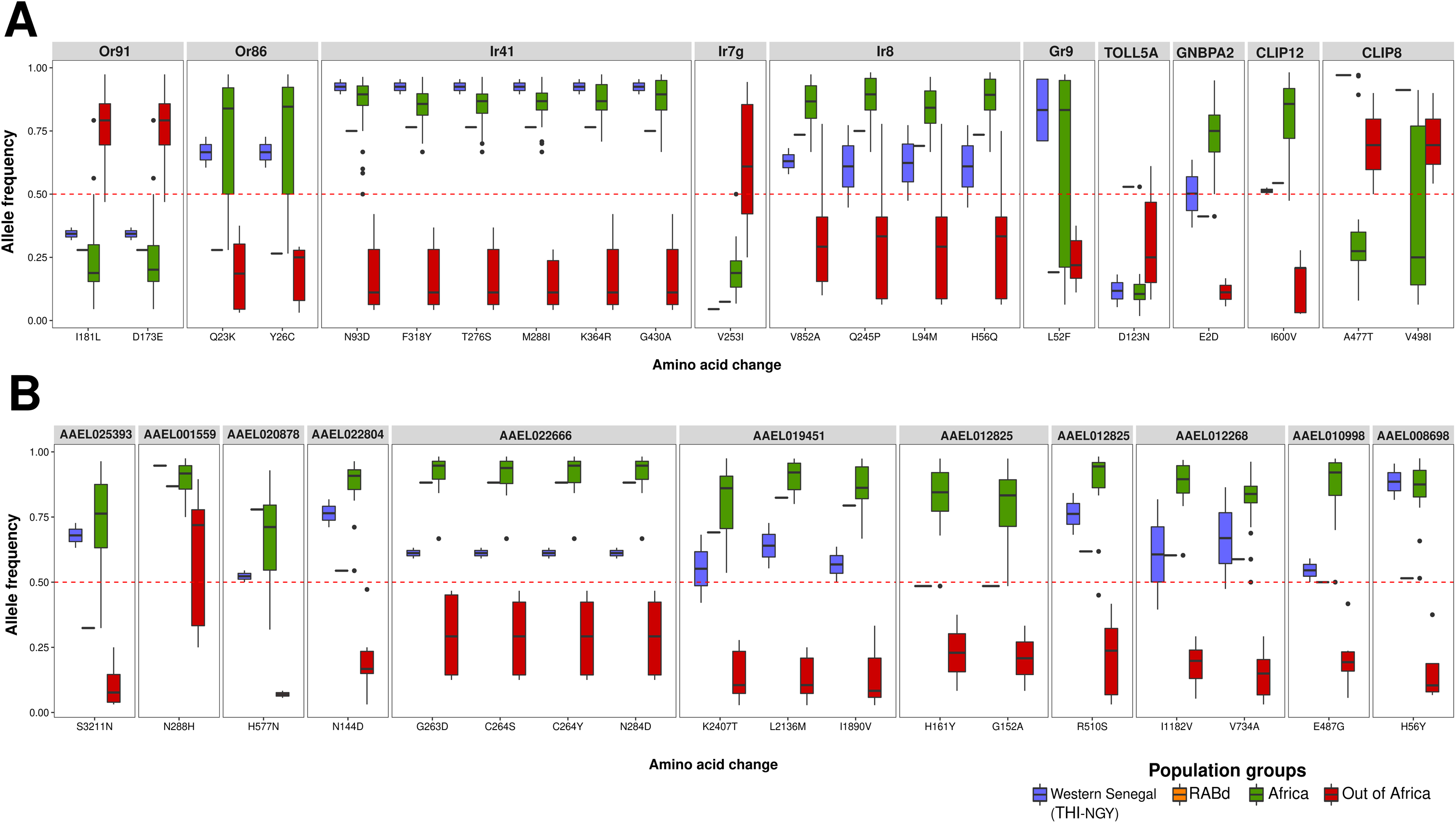
Molecular markers differentiating Aaa and Aaf mosquitoes. Box plots depicting the alleles frequencies (on the Y-axis) of nonsynonymous mutations depicted as resulting amino acid change (on X-axis) within selected “*Aaa molecular signature*” genes. Populations are divided and color-coded in four groups, one for all out-of-Africa mosquitoes and three groups for African mosquitoes: Western Senegal (THI-NGY), RABd, Africa (others). (**A**) Examples of genes with known function in chemosensation (Or91, Or86, Ir41, Ir7g, Ir8, Gr9) and immunity (Toll5A, GNBPA2; CLIP12, CLIP8) are shown in separate panels. (**B**) Examples of genes with unknown functions (AAEL025393, AAEL001559, AAEL020878, AAEL022804, AAEL022666, AAEL019451, AAEL012825, AAEL012783, AAEL012268, AAEL010998, AAEL008698). The red dotted line shows the middle value (0.50) of the allele frequency in which all values are distributed.

### Genes associated with diverse neuronal functions are positively selected in out-of-Africa mosquitoes

We further estimated selective constraints by calculating the ratio of non-synonymous (Ka) to synonymous (Ks) substitutions (dN/dS ratio) for the 13,503 *Ae. aegypti* genes that mapped SNPs in their protein-coding exons. We found 12,991 genes evolving under negative selection, which are mostly shared across all tested populations, with none of the five clusters identified offering an unambiguous distinction between Aaf and Aaa mosquitoes. By contrast, we found that most local adaptation across Aaa populations seems to be driven by 1,033 genes (from 9 clusters) out of the 5,242 genes that were found evolving under positive selection. We consider this set of 1,033 genes as “*Aaa molecular signature*” because they clearly differentiate African from out-of-Africa populations (**Fig. 5**). The “*Aaa molecular signature*” genes were distributed across the three chromosomes, encompassing the regions from 128.3 to 287.7 Mb and 284.5 and 344.8 Mb on chromosome 2, which harbor Quantitative Trait Loci (QTLs) previously linked to higher vector competence for Zika virus in mosquitoes from Guadeloupe vs Gabon (5). Chemosensory and immunity genes were the most represented ones among the “*Aaa molecular signature*” genes (**Fig. 5**). We detected several olfactory-associated genes evolving under positive selection across several out-of-Africa populations, including *Gr1* in JED and BNK; *Gr34* in JED and SAS; *Gr20* in SAM, TAP, SAS, and BNK; *Obp*jj7a in JED, SAS, and BNK; *Obp14* in JED; *Or13* in JED and SAS; *Or44* in TAP and SAS; *Or45* in JED, SAS and BNK; and *Or4* in TAP and JED. Across tested mosquitoes from Asia and/or America, signals of positive selection were found to be mostly local, with concordant results from the PCA-outlier analysis in *Gr1, Gr8, Gr20a, Ir7g, CLIB1* and *GPXH2*; the cytochrome P450 CYP6Z7, which has been frequently associated with resistance to pyrethroids (37–39), showed pervasive positive selection, showed pervasive positive selection in remarkable contrast to most negatively selected P450 genes, further underscoring its physiological role and the significance of these results.

**Fig. 5.**
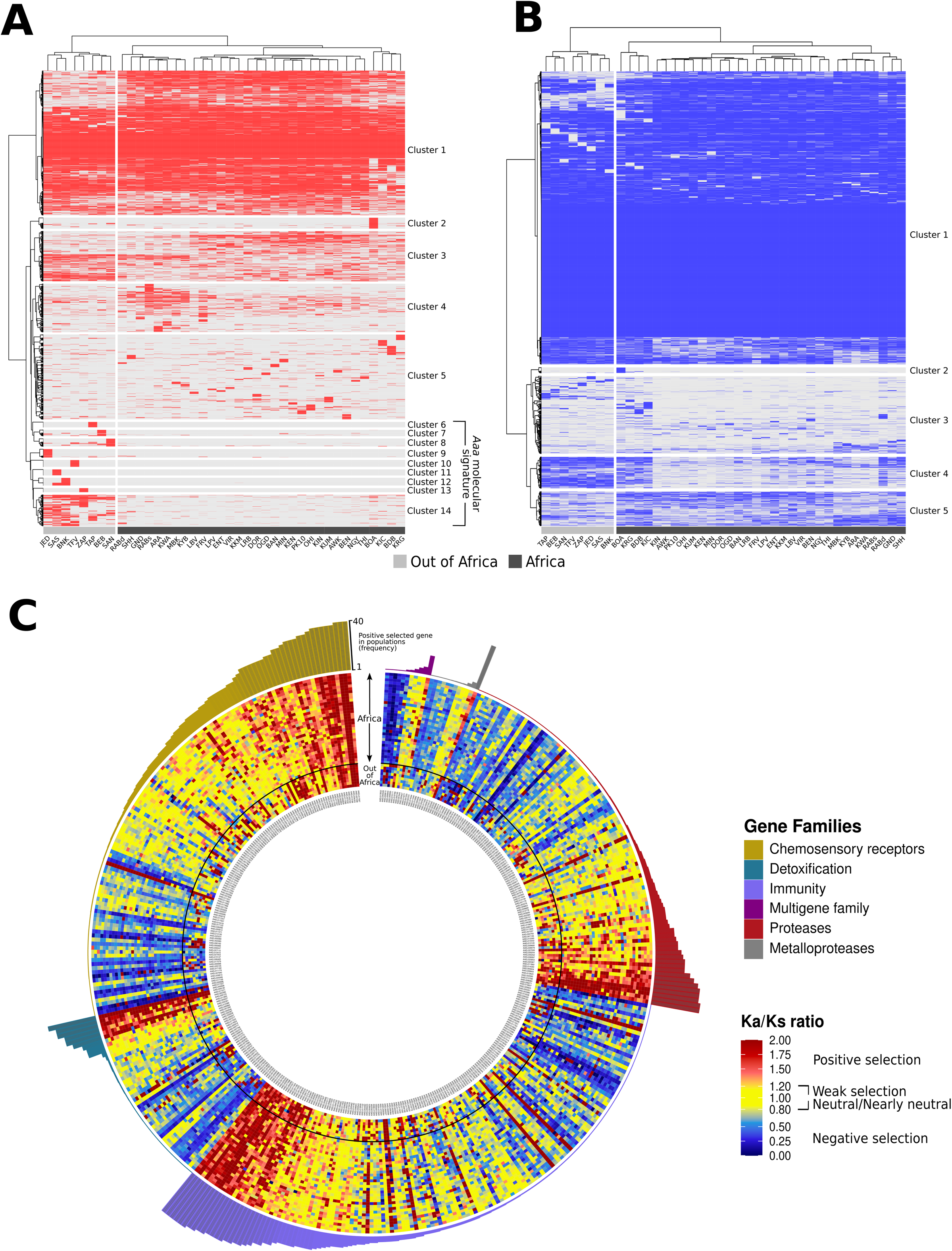
Identification of protein-coding selection and *“Aaa molecular signature”* from >13,500 genes in *Ae. aegypti*. The heatmaps in the top panels show the clustering of genes under positive (A) or negative (B) selection in at least one population. Ka/Ks ratio values of A and B were transformed in a binary matrix based on the Ka/Ks thresholds (see below) defining the *presence* or *absence* of selection acting on genes (on the Y-axis) across Aaf and Aaa populations (on the X-axis), separately. (**A**) When Ka is much greater than Ks (*i.e.*, Ka/Ks >>1) (56), we found 5,242 genes as evolving under positive selection, *i.e.*, their variants are likely beneficial and being promoted. (**B**) When Ka is much less than Ks (*i.e.*, Ka/Ks << 1) (56), then 12,991 genes were estimated as evolving under negative selection, *i.e.*, their variants are likely deleterious and being eliminated. (**C**) The central circle of this polar plot shows the distribution of 419 genes evolving under positive selection (in red-color scale) in at least one population as selected from a catalogue of 1,132 genes belonging to one of the six major functional categories considered relevant for domestication in *Ae. aegypti*. All 40 populations analyzed, 32 from Africa and 8 from out-of-Africa, are located over the Y-axis, while the 419 genes are located over the X-axis. The outer circle shows the cumulative frequency (in bars) of each gene selected positively across the 40 populations, according to the functional category it belongs to as in the legend.

Among the “*Aaa molecular signature*” genes is notable the presence of genes associated with neuronal and hormonal functions such as the N-methyl-D-aspartate (NMDA) receptor (AAEL008587), which belongs to the family of ionotropic glutamate receptors playing a major signaling role in the central nervous system and in neuromuscular junctions (40); the *capa* gene (AAEL005444) belongs to the PK/PBAN family of neuropeptides, which in *Drosophila melanogaster* encodes for pyrokinin-1 and periviscerokinins (41), and has been shown to regulate diverse physiological functions ranging from feeding behavior (42) to the regulation of water balance (43); tomosyn (AAEL006948), a motoneuron receptor that influences homeostatic synaptic plasticity (44); the synaptic vesicle protein synaptotagmin (AAEL001167) (45), and several G-protein coupled receptors (GPRs), a category of proteins enriched among *Anopheles gambiae* brain peptides (46).

### Endogenous Viral Elements contribute to differentiate out-of-Africa and African mosquitoes

Recent experimental evidence expands the immunity toolkit of *Ae. aegypti* to nonretroviral endogenous viral elements (nrEVEs), which interplay with the piwi-interacting (pi) RNA pathway to control cognate viral infections (47,48). The genome of *Ae. aegypti* harbors 252 nrEVEs (hereafter *reference* nrEVEs), > 50% of which are at least 4 MYA being shared with *Ae. mascarensis* (47,49). *Reference* nrEVEs coexist with 64 “*new”* nrEVEs, which are exclusively detected in wild mosquitoes, and their distribution significantly differed in African and out-of-Africa mosquitoes. A total of 5 *new* nrEVEs (Guato_2, CFAV_EVE-4, CFAV_EVE_1, CFAV-EVE-5, *Aedes aegypti* toti_like-7) are unique of out-of-Africa mosquitoes and, overall, *new* nrEVEs have a higher nucleotide identity to cognate viruses than *reference* nrEVEs. With the exception of three integrations from the *Liao Ning* virus belonging to the *Seadornavirus* genus (Reoviridae family), which includes emerging pathogens (50), all *new* nrEVEs are from limited number of ISVs. We also observed frequent rearrangement events following integration, especially for nrEVEs with similarity to flaviviruses that map in piRNA clusters, supporting the conclusion that nrEVEs contribute to the genetic flexibility of *Ae. aegypti* piRNA clusters.

## DISCUSSION

Domestication is a complex, multifactorial and species-specific process traditionally associated with vertebrates, among animals. Still, domestication events have also occurred in a few invertebrates such as shrimps (*Panaeus* spp.), the silk moth *Bombyx mori* and *Ae. aegypti* (1,51,52). In the case of *Ae. aegypti*, it took only a few hundred years for the species to become globally invasive and human specialist (11,12). *Aedes aegypti* domestication process has resulted in changes on different aspects of its bionomics (e.g. vector competence, reproductive behavior and host feeding preferences) and, by consequence of human interventions, in insecticide tolerance in just a few thousand years (6,12,24).

This work represents the most comprehensive effort to date to identify genomic variants and footprints of genomic selection tracing the evolutionary divergence and behavioral switch of *Ae. aegypti* populations to domestication. To revisit this matter with unprecedented phylogenetic resolution, we used the high-quality reference assembly AagL5 (32) and the genomes of 511 African and 123 out-of-Africa mosquitoes that were sampled and sequenced from 14 countries across four continents. From >300 million high-confidence SNPs detected throughout the whole *Ae. aegypti* genome, we found protein-coding variants that can significantly differentiate Aaa from Aaf mosquitoes. We call this group of genes “*Aaa molecular signature*” genes. A total of 236 “*Aaa molecular signature*” genes harbor nonsynonymous mutations that occur at statistically different frequencies between Aaa and Aaf mosquitoes. Thus, they could be used to distinguish the two *Ae. aegypti* subspecies molecularly. In the following, we discuss the population structure context under which these “*Aaa molecular signature*” genes were identified, highlighting their association with expected (olfaction) and novel functional hallmarks of domesticated behaviors in *Ae. aegypti*.

Our genetic structure and phylogenetic analyses performed over 1.5 million biallelic and non-redundant SNPs, led us to confirm a major genetic divergence between African and out-of-Africa populations, with the former being clearly genetically structured into Western, Western/Central, Central and Eastern metapopulations. These results are strongly supported by the pairwise-Fst genetic distances obtained across all populations, with the unique exception of RABd. We consistently found that RABd formed a cluster, which separated from other African populations and was phylogenetically more closely related to, and shared the lowest genetic divergence with, JED, in comparison to all other tested populations. Empirical observations of closer relatedness between Aaa populations and behaviorally divergent ‘‘domestic’’ and ‘‘forest’’ mosquitoes of Rabai have been reported previously, with contradicting hypotheses concerning the origin of such ‘domesticated behavior’ (6,10,16,53,54). The findings of our study provide compelling evidence for a “back to Africa” event, indicating a recent reintroduction from Saudi Arabia into Kenya of Aaa mosquitoes, which remained localized as indicated by extensive inbreeding.

Our admixture and PCA analyses also provide strong support for the separated genomic clustering of the human-feeding mosquitoes sampled from the Senegalese NGY and THI populations (6). Results from ML-phylogenies further showed that NGY and THI are closely related to out-of-Africa mosquitoes. Importantly, not only the branch grouping both THI and NGY with out-of-Africa populations is basal to all African population, but it also supports a high genetic divergence between NGY/THI and out-of-Africa populations when compared to other African populations, which is consistently confirmed by all pairwise-Fst distances. We also demonstrate that the close phylogenetic relationship of both THI and NGY with Aaa populations is not the product of admixture events, given that the F3 test was rejected in all cases. F3 results discarding admixture with out-of-Africa populations extend to all tested African mosquitoes. Altogether, these findings are consistent with inferring that NGY and THI derive from an ancestral domesticated population, rather than representing recent re-introductions and/or admixture events between African and out-of-Africa mosquitoes (9,11,13–15,18,55).

The genome-wide SNP divergence between African and out-of-Africa populations, which agrees with previous studies (6,13,16,18), is further endorsed by the differential clustering of nrEVEs between the two subspecies, with 5 nrEVEs being exclusive of out-of-Africa populations. These Aaa-specific nrEVEs belong to a group of 64 novel, and PCR-validated, nrEVEs that we strongly suggest being the outcome of recent integration events given their higher nucleotide identity to cognate viruses. The population structure that we found, meaning a unique origin for all out-of-Africa domesticated mosquitoes with NGY and THI representing ancestral domesticated mosquitoes and no current admixture between African and out-of-Africa mosquitoes or new introductions of Aaa into Africa, except for RABd, was the basis to continue in the identification of genomic variants and footprints of genomic selection between out-of-Africa domesticated and generalist African mosquitoes. To test whether or not switches to long enduring domesticated behaviors in *Aedes aegypti* have a strong and multi-loci genomic basis, we joined two well-known approaches: first, we identified genomic variants more strongly associated in all populations of the two subspecies than expected only by genetic drift; second, we identified all those positively selected protein-coding variants that can unambiguously differentiate patterns of adaptation in African versus out-of-African populations. We found that the genetic divergence between Aaa and Aaf is highest in a group of genes from the prediction of both approaches, that we call “*Aaa molecular signature*” genes. Given that results from these estimations are sensitive to the number of populations that are tested and their grouping, we performed our analyses on all populations and statistically validated population divergence prior to calling outlier SNPs. While some “*Aaa molecular signature*” genes may reflect local adaptation independently of domestication, in gene families associated with functional redundancy, local adaptation signals may still be domestication signals.

Our candidate “*Aaa molecular signature*” genes include olfactory genes, which mediates sensing of volatiles, a function used by *Ae. aegypti* females to locate both a host for blood feeding and a breeding site for oviposition (58,59). Olfaction is governed by multi-gene families and has a highly redundant organization in *Ae. aegypti*, with multiple receptors in the same neuron and individual variability, which is different from the canonical organization “one receptor-one neuron-one glomerulus” observed in *Drosophila melanogaster* (58). This level of redundancy increases the breath and the flexibility of volatile perception, and it may be linked to local adaptation at the genomic level. Despite this redundant organization, a few candidate receptors have been identified as having a major role because of their either ubiquitous co-receptor function (i.e., *Orco*, *Ir8a*, *Ir25a,* and *Ir76b*) or prevalent sensing of human-relevant compounds, such as sulcatone by *Or4*, CO_2_ by *Gr3*, and lactic acid by *Ir8a* (26,60–63). We identified pervasive negative selection in *Orco* and *Gr3*, underscoring their physiological role. Consistent with the hypothesis that functional redundancy may entail local adaptation, we detected strong genomic signals of local adaptation across several olfactory-associated genes such as *Gr1* in RABd and all out of Africa; *Gr20* in Kenyan populations (MBK, ARA, GND, KYB, KWA, SHH, and RABd, RABs) and NGY; O*r26* in THI, NGY, RABd, RABs; *Or44* in TAP and SAS; *Or45* in JED, SAS and BNK; *Or86* in RABd and out of Africa; *Or4* in TAP and JED; *Gr8* and *Gr20a* in Kenyan populations (MBK, ARA, GND, KYB, KWA, SHH, and RABd, RABs) and NGY; *Ir7g* in THI, NGY, both RAB, and out of Africa populations*. Or4* is particularly significant as different *Or4* alleles are known to circulate across *Ae. aegypti* populations, with levels of *Or4* expression being strongly predictive of preferential attraction to humans (26). Here, we circumscribed our analyses to coding sequences, thus we are possibly missing variants that could regulate *Or4* expression. Such analyses would require complementary transcriptomic data assessing gene expression and extensive functional validation, which are beyond the scope of our current work. In the co-receptor *Ir8a*, which is expressed specifically in antennal neurons and is required for perception of lactic acid, a component of human sweat (60), we identified nonsynonymous mutations occurring at significantly different frequency in Aaa and Aaf mosquitoes. In NGY and THI, which are African mosquitoes that behave like Aaa in their preference for humans (6), these mutations appear at frequencies which are intermediate between those detected in Aaa and Aaf mosquitoes, underscoring their significance and pointing to functionally assessable targets for future endeavor.

As expected, our list of “*Aaa molecular signature*” genes include additional genes belonging to major gene families such as detoxification, immunity, and proteases, which have been shown to impact host seeking behavior, vector competence and overall response to external stimuli (6,32,64,65). Notable examples include *Toll5A,* which interacts specifically with Spaetzle1C resulting in regulation of immunity and, primarily, fatty acid metabolism (66,67), as well as several clip-domain serine proteases, which are upregulated following *Ae. aegypti* infection with different pathogens (68,69). Although >50% of “*Aaa molecular signature*” genes lack a functional annotation, also notable is the presence of genes associated with broad hormonal and neuronal functions such as the NMDA receptor, the *capa* gene, tomosyn and synaptotagmin (41–43,45), suggesting that the behavioral shift to domestication may rely on the fine regulation of metabolic and neuronal functions, more than the role of a few major genes. In support to this hypothesis, domestication in rabbits, which occurred rapidly in the past 1500 years, resulted in a shift in primarily SNPs nearby genes associated with brain and neuronal development (1,70). Additionally, a mutation affecting the *thyroid-stimulating hormone receptor*, which controls photoperiodic diapause and reproduction, is widespread but not fixed in domestic chickens (71,72), and candidate domestication genes in the silkworm (*Bombyx*) include genes involved in energy metabolism, reproduction and silk gland activity (69).

Altogether, our findings robustly suggest that domesticated behaviors in *Ae. aegypti* have evolved by shifts in allele frequencies and codon selection at many loci, owing to diverse selective pressures caused by local adaptations to microgeographic and anthropogenic changes worldwide. In particular, selection on olfactory genes and their related nonsynonymous variants, which are expected to be relevant for generating the behavioral switch in *Ae. aegypti* to domestication, are found to be strongly influenced by genetic background and population history. The genetic diversity richness of the generalist African populations found in both repetitive and non-repetitive regions throughout the whole genome strongly suggests that retention of ancestral polymorphisms is likely the main genetic source for the evolution of complex evolutionary dynamics in the domesticated behaviors of *Ae. aegypti*. The absence of introgression between the Aaa and Aaf populations analyzed here, and the 2-fold reduction of SNPs, nucleotide diversity, rare variants, and high genetic drift that we found across out-of-Africa populations strongly endorse this hypothesis, although other genomic events, recent retention of polymorphisms due to local introgressions and convergent evolution on certain loci are not to be discarded (*e.g.*, (73)). Retention of ancestral allelic variants based on microsatellite markers was suspected to occur in *Ae. aegypti* (13,18,20), but was only reported in other human-feeding mosquitoes recently, such as *Anopheles gambiae* (74–77), *Culex nigripalpus* (78), and *Cx. quinquefasciatus* (79). Particularly notable is the presence of alternative allelic variants at low frequency (*i.e.*, “standing variation”) in the same protein-coding genes that we found to be evolving under nearly neutral or weak (positive/negative) selection across most African populations (**Fig. 5c**), which may be maintained for longer periods of time beyond neutral expectations (80). This is the first large-scale observation of selection over preexisting standing variation in *Ae. aegypti*, a phenomenon that has been also reported in *Daphnia* (79) and a few other organisms (82–85). By selecting from a rich stock of ancestral and weakly evolving standing variants from African populations, we observe that some of the *“Aaa molecular signature”* genes such as the odorant receptor gene AAEL000616 found within modules linked to active periods or sleep-like states (86), the chymotrypsin JHA15 (AAEL001703) found to be highly expressed before a blood meal (87), and the immune-related recognition gene AAEL019958 found to be exclusively expressed in ovaries of female mosquitoes (88), and several chemosensory genes have switched to directional selection in several out-of-Africa populations, where the fast emergence of novel adaptations can be easily promoted by the redundant organization of olfaction to cope with novel geographical and anthropogenic evolutionary pressures.

Relative to recent efforts in other human feeding mosquitoes, our work provides a unique resource for the study of population genetic structure and genome selection on a microgeographic scale for one of the fastest evolving arbovirus vectors worldwide. We also pinpoint to further candidate genes involved in neuronal functions as targets of genetic manipulation and neurogenetic approaches recently developed for mosquitoes (58,89–92).

## Methods

### Mosquito samples

We studied the whole genome sequence of 694 *Aedes* spp. mosquitoes. This sample size included previously published WGS data of *Ae. aegypti*, *Ae. mascarensis* and *Aedes albopictus* (6,10,47,93) and WGS data of 97 *Aedes* spp. mosquitoes that we processed from Burkina Faso, Ethiopia, Brazil, Saudi Arabia, Russia, Cameroon and New Caledonia. With the exception of New Caledonia from where we received eggs through the ‘Infravec2’ project (https://infravec2.eu/), from all other sites we received adult mosquitoes preserved in ethanol 70%; these mosquitoes had been sampled either as larvae from tires, backhoe buckets and various surrounding larval habitats or as adults through ‘BG-sentinel’ traps. Mosquitoes from Cameroon are from a colony established from eggs collected in Bénoué; females were sampled at the 12th generation after colony establishment. Genomic DNA was extracted from individual mosquitoes using the Wizard Genomic DNA Purification Kit, according to the manufacturer’s protocol, at the University of Pavia for all specimens, apart from mosquitoes from Brazil, which were processed *in loco.* Genomic DNA was sent to Macrogen for individual DNA library preparation with TruSeq DNA PCR-free reagents and sequencing to a minimum of 20X coverage (24X on average) in paired-end 150 bp reads with the Illumina HiSeq X Ten platform. FASTQ files of all WGS datasets were subjected to quality control by using FASTQC v0.11.9 (94). Sequencing data were deposited to the NCBI SRA under the accession BioProject ID: PRJNA943178.

### Alignment to the reference genomes

Raw reads of each of the 694 WGS datasets were trimmed using Trimmomatic v0.39 (95), then the 21 *Ae. albopictus* WGS data were aligned to the *Ae. albopictus* Foshan FPA genome assembly (96), the remaining WGS data were aligned to the current *Ae. aegypti* reference genome assembly, AaegL5 (32); both assemblies were downloaded from VectorBase (https://vectorbase.org/). The BWA MEM algorithm v0.7.17.r1188 was used for all alignments (97). For each sample, genome mapping statistics were calculated with Qualimap v2.0 (98) and alignment quality statistics were obtained with bamtools (99). For WGS data mapped to the *Ae. aegypti* genome, gene coverage was calculated for the 14,677 genes reported in AaegL5 with the program mosdepth v0.2.9 (100). For a total of 35 samples, less than 50% of the reads aligned to AagL5. To clarify this result, we reconstructed the sequences of the ribosomal Internal Transcribed Spacer 1 and 2 (ITS1 and ITS2) of these 35 sampled by using the *LT_finder* script of the ViR pipeline (49) and the 5.8S rRNA sequence of *Ae. aegypti* (accession M95126, region coordinates 574-755 in AagL5) as reference. The reconstructed ITS1-ITS2 sequences were then blasted, with *blastn*, against the ‘nt database’ of NCBI with default parameters through the BLAST webserver (https://blast.ncbi.nlm.nih.gov/) to confirm species identity. All samples from Russia and Ethiopia showed the highest identity to ITS sequences of *Ae. albopictus* (identity>97.24%, 100% query coverage, and E-value of 0) or *Ae. simpsoni* (identity>97.66%, 100% query coverage, and E-value of 0), respectively. WGS data from 9 additional mosquitoes had low (≤2 reads) and/or partial coverage (≤45%) to AaegL5, but their ITS1-ITS2 reconstructed sequence did not yield results when searched against the ‘nt database’ of NCBI, suggesting either uncharacterized species or sequencing quality issues. To avoid biases in the analysis, these WGS datasets were excluded resulting in final dataset of 634 mosquito genomes from 39 populations, for which ≥96% of reads mapped to AagL5. In these WGS samples, 95% of the 14,677 *Ae. aegypti* genes were covered with ≥5 reads; the remaining 5% of genes, which were covered with ≤4 reads, mapped in *contigs* that have not been assigned to any chromosome yet.

Females were identified among these 634 mosquitoes by the complete absence of coverage on *Nix* (AAEL022912); males are expected to have coverage over the protein coding region of *Nix* (≥1 read) (101), which was estimated with Samtools v1.4 (102). In presence of coverage over *Nix,* we also verified coverage of the *myo-sex* gene (AAEL021838) to further support the sex association to males. To verify amplification of the *Nix* gene from sperms stored in female spermathecae, we sampled males, virgin females and females collected after copulation. DNA of each of these samples was extracted with the Wizard Genomic DNA Purification Kit (Promega) following manufacturer recommendation. DNA was amplified with a nested PCR using in the first PCR primers Nix_aeg_PCR-F: 5’-ACGGAAGAGCGAATTGCACA and Nix_aeg_PCR-R: 5’-GTCAAACCGTCTGAGCGTCT and in the second reaction primers Nix_aeg_nPCR-F: 5’-AGCGTGCTTCAGAATAATTACGG and Nix_aeg_nPCR-R: 5’-GTTTTGATGCGGTGAGTGCC. PCR reactions were assembled using the DreamTaq Green PCR Master Mix (Thermo Scientific) following manufacturer’s instructions. 1 uL of DNA extract was added to reach a final volume of 25 uL. PCR reactions were performed in a thermal cycler (Eppendorf™ Mastercycler Nexus Gradient) with, after an initial denaturation for 3 minutes, 35 cycles at 95°C for 30 s, 52.4°C or 53.3°C for 30 s for the first or second PCR, respectively, and an extension of 25 s at 72°C, followed by a final extension for 10 minutes at 72°C. PCR products were visualized using a Bio-Rad Gel Doc TM EZ Imager following electrophoresis in a 2% (w/v) agarose gel.

### Recalibration of alignments and variant discovery

The 634 WGS data mapped to the *Ae. aegypti* genome (AagL5 assembly) were further processed following the best practices recommendations from the Genome Analysis Toolkit (GATK) (103,104). First, the program Picard v2.23.0 (Broad Institute, 2019; https://broadinstitute.github.io/picard/) was used to sort aligned reads and mask optical duplicates. Then, local realignments were performed with the GATK package v3.81.08 (105) over regions mainly characterized by indels (insertions and deletions) and read mate coordinates of realigned reads were recalculated with the Picard program. Finally, the Base Quality Score Recalibration (BQSR) was calculated for each alignment with the GATK package. To improve alignment during the recalibration step, we provided to GATK a set of known indels and Single Nucleotide Polymorphisms (SNPs). This set of variants was built with two different approaches: 1) known SNPs from literature and 2) *de novo* SNPs estimates for our sequenced mosquitoes through bioinformatic analyses. Both procedures are described next.

#### SNPs collected from literature

An initial collection of 485,431 SNPs was obtained in a variant calling format (vcf) file from nine published studies (23,106–113). From this collection, 112,969 SNPs (23.27%) have experimental support such as high-throughput genotyping chip ‘Axiom_aegypti1’ (106–110), while the remaining 372,462 SNPs (76.73%) were estimated only through bioinformatic analyses (23,111–113). The genomic coordinates of all nine SNPs datasets were re-mapped from the *Ae. aegypti* assembly AaegL3 to the AaegL5 version through the lift over strategy by the VectorBase website. A SNP dataset originated from exome analysis (13) was also added to our study. For this dataset, we performed a local lift over with the *Flo* pipeline to re-map all genomic coordinates from the AaegL1 to the AaegL5 assembly (114). We used the UCSC chain files (113), the gnometools package (114), and the pblat-cluster v36×2 (117,118) to accelerate the sequence search mapping across genome assembly versions. Finally, we used Crossmap (119) to perform the genomic conversion of coordinates for the set of known SNPs over AaegL5. Overall, after merging sites and removing overlapped redundant variants, from these 10 SNP datasets a total of 304,428 SNPs were mapped across 10,185 *Ae. aegypti* genes and 278 contigs annotated in AaegL5.

#### De novo SNP discovery

Two variant callers, GATK v3.8.1.0 (105) and Freebayes v1.3.1 (https://github.com/freebayes/freebayes) were applied to identify SNPs from each of the 634 WGS samples mapped to AagL5 to improve SNP prediction because both variant calling tools have a high false discovery rate (FDR) when compared to a true set of known SNPs (120–122) and both tools are known for scoring best at one, rather than in all, quality parameter measurements (120,123). We further implemented a strict filtering protocol for each set of identified variants to complement the power prediction of both variant calling tools and increase specificity and sensitivity. First, variant caller predictions were performed with GATK and Freebayes for each sample, separately.

Then, raw SNPs and indels calculated from Freebayes were extracted and filtered with: 1) a minimum deep coverage of 5 reads (DP≥5); 2) a minimum mapping quality of 20 (MQ≥20); 3) a minimum base quality of 20 (QUAL≥20) and 4) a minimum allele frequency ≥ 1% (maf≥0.01), as calculated with BCFtools (102). Filtering of the GATK called variants was implemented with the GATK protocol for SNPS: QUAL < 30.0, QD (QualByDepth) < 2.0, FS (FisherStrand) > 60.0, MQ (RMSMappingQuality) < 40.0, SOR (StrandOddsRatio) > 3.0, MQRankSum (MappingQualityRankSumTest) < −12.5, ReadPosRankSum < −8.0; for INDELs: QUAL < 30.0, QD < 2.0, FS > 200.0, ReadPosRankSum < −20.0). All SNPs with a close proximity of 10 base pairs to an indel and with a strong strand bias (p<0.001) were removed with BCFtools. Finally, we merged all sites and removed all overlapped redundant variants from the known and *de novo* approaches, which generated a final high-quality collection of 82,686,298 indels and 207,724,254 SNPs that we used in the recalibration step, as described above.

A refined variant caller prediction was performed only with the GATK protocol for all recalibrated alignments in each of the 39 populations, separately. Raw SNPs and indels were extracted and filtered with the same filtering parameters using GATK, as described previously. After this last filtering process, a total of 314,365,358 SNPs were obtained as the core dataset of our analyses, while indels were not considered in further analyses.

### Pairwise relatedness of individuals in *Ae. aegypti* populations

We evaluated the *degree of relatedness* among the 634 individuals within each of the 39 populations and removed closely related individuals (*i.e.*, full siblings) to avoid any bias. To this end, we first removed all SNPs detected over repetitive regions of the *Ae. aegypti* genome by using the genomic coordinates reported by (32); and then we extracted all biallelic SNPs present in at least 90% of individuals within each population. Next, highly linked loci were eliminated using the function *snpgdsLDpruning* (ld.threshold=0.01) of the SNPRelated R package (124), and the corresponding matrix of ‘relatedness coefficients’ (*k*) was generated with the Identity-by-Descent (IBD) measurement based on maximum likelihood estimation (MLE) by using the function *snpgdsIBDMLE* included in the same R package. Highly genetically related individuals were classified and filtered in each population using two conservative cutoffs for African and out-of-Africa populations, as previously suggested by (6): *first cousin and closer relationships* (*k≤*0.05) for African populations and *siblings* (*k*≤0.20) for out-of-Africa populations. We repeated this analysis by using a dataset of 1,000,000 randomly sampled SNPs across the whole genome per population, and manually compared both approaches by identifying the individuals, as well as the total number of individuals to be removed. Through this process, we removed 95 individuals, leaving a dataset of 539 individuals. Among these 95 related individuals, 15 corresponded to the previously called ‘domesticated’ mosquitoes of the Rabai population (6). To confirm the *relatedness groups* estimated with our protocol, an analysis on the covariance of genotypes was performed with the *pca* function of the plink v2.0 package. Based on this analysis, all domestic individuals from Rabai were reintroduced to our dataset, resulting in final set of 554 *Ae. aegypti* genomes for further analyses.

### Distribution of SNPs and genetic diversity along the *Ae. aegypti* genome

Unless otherwise stated for all further analyses, we used *Ae. aegypti* genomic coordinates as reported in AaegL5 (32). We mapped the entire set of 314,365,358 filtered biallelic and multiallelic *Ae. aegypti* SNPs to compare the distribution of SNPs across each centromeric region and chromosome arms (*1q*, *1p*, *2q*, *2p*, *3q*, *3p*) and used a paired t-test to find significant difference between and among chromosome arms in African and out-of-Africa populations. We further estimated the total number of SNPs counts in chromosomes (hereafter ‘chromosomal SNPs’) or contigs (hereafter ‘unassigned SNPs’), and, in each of these regions, we counted SNPs in exons, protein coding regions (CoDing Sequences, CDS) and untranslated (5’-UTR and 3’-UTR) regions and further split SNPs based on whether they occurred in repetitive (R-SNPs) or non-repetitive regions (NR-SNPs) by using the function SelectVariants, options *intervals/excludeIntervals,* in GATK. R-SNPs counts are listed for Transposable Elements (TE), low complexity sequences and unclassified repeats.

Focusing on NR-SNPs, we measured genetic variance in terms of SNP density, genetic diversity (π) and Tajima’s D using the VCFtools package (125). A genome-wide scan with different non-overlapping sliding windows (10kb, 50 kb, 100 kb, 250 kb, and 500 kb) was performed to calculate both basic statistical descriptors for genetic variation. Finally, we identified the presence of ‘SNP singletons’ (*i.e.*, a SNP present in one single individual of a population) with the VCFtools package (options *singletons* and *positions*) and estimated their counts and distribution across populations using a custom R script.

### Population Genetics Analyses

Chromosomal biallelic SNPs that were found in at least 80% of all individuals were extracted and further filtered out using plink v2.0, to avoid sampling genotyping errors (126), if highly linked (option indep-pairwise: window size=50, step size=10, and R^2^=0.1), having a MAF<0.01 or showing a significant deviation from Hardy-Weinberg equilibrium (HWE) (p<0.001). This procedure led to a total of 1,530,512 SNPs that were used to assess the genetic relationships across populations using the *pca* function of the plink v2.0 package, for admixture analysis using the ADMIXTURE software v1.3.0 (127), and to estimate F_ST_ population scores using VCFtools (125). As described in (128), we ran ADMIXTURE on all individuals and varied the number of genetic clusters (K) from 1 to 39 to identity the number of clusters that minimizes the cross-validation error. We performed the PCA and admixture analyses on different genomic scales (*i.e.*, WG, R, NR and exon regions) to test for differences in the distribution of genetic variation across the mosquito genome that might potentially induce distinct effects on the populations structure. For exon regions, 1,000 bootstrap replicates for every dataset with a cluster (*k*) value from 2 to 39 were carried out to further support the identification of the optimal cluster number. A matrix of all-vs-all pairwise comparisons of the F_ST_ population scores was built using VCFtools and a custom PERL script to estimate the genetic divergence across populations. All populations were then grouped according to a *complete* hierarchical clustering by using an *euclidean* distance and 1,000 bootstrap replicates with the pvclust R package (129).

We further investigated the phylogenetic relationships among the 554 *Ae. aegypti* genomes by building a ML phylogenetic tree with all SNPs from exons of protein-coding genes that were present in 100% of individuals across all populations (named here as “core-exome SNPs); we define this phylogeny as “tree of individuals”. This set of SNPs was transformed into a phylip format with the vcf2phylip program (https://github.com/edgardomortiz/vcf2phylip/blob/master/vcf2phylip.py) and the corresponding phylogeny was reconstructed with a GTR+CAT substitution model (-m ASC_GTRCAT) that includes an ascertainment bias correction for SNPs (ass-corr=lewis); the statistical robustness of the phylogeny was assessed with 1,000 bootstrap replicates using RaxML v8.2.12. Using the same set of SNPs, we also derived a ML phylogenetic tree based on SNP frequencies estimated within each population; we define this phylogeny as “population tree”. This tree was built with the TreeMix program after 1,000 bootstrap resampling of the dataset (33). For both phylogenetic trees, *Ae. albopictus* was used as an outgroup. The F3 statistics implemented in the TreeMix program (program *threepop*), which measures the covariance of the differences in allele frequencies among three populations, was used to test the genetic admixture of THI and NGY populations with respect to out-of-Africa populations. The association test of these populations is depicted in a tree topology of the type (A,B;C), where *C* is either THI or NGY, and *A* and *B* represent all possible combinations of the out-of-Africa populations. Genetic admixture was established based on z-scores as a test statistic and a conservative threshold (z-score*≤*-3.0). Similarly, we performed the same type of analysis to all-vs-all African populations, particularly focused in recent populations that have shown human seeking behavior THI, NGY, OGD, KUM (6,130).

### Outlier analysis and local adaptation in populations of *Ae. aegypti*

We screened for SNPs showing unusual patterns of genetic variation along the *Ae. aegypti* genome, with the hypothesis that they contribute to the genetic differentiation among populations. To this end, we used the “outlier method” implemented in the PCAdapt R package v4.3.3 (35), which calculates a PCA from SNP data to search for loci that are atypically related to the population structure by decomposing the ‘total’ genetic variation into ‘axes’ of genetic variation (*K*) called “principal components” (PCs). PCAdapt further calculates the correlations between SNPs and a specific number (*K*) of retained PCs, so that SNPs showing an excessive relation with the population structure are defined as outliers and suggested to be candidates for local adaptation. We performed this analysis using the 1,530,512 biallelic chromosomal SNPs that were detected in at least 80% of individuals across all populations; this analysis was performed separately for each chromosome. Importantly, we did not impose any population clustering, but estimated the ‘optimal *K* axis’ running PCAdapt with an excess of K=20 and assigning the ‘optimal *K’* based on three different approaches: 1) the Cattell’s rule, which keeps PCs that correspond to eigenvalues to the left of the lower straight line in the screeplot (131); 2) the Tracy-Widow test (p-value<0.05), which was applied to the eigenvalues by using the program *twstats* from the EIGENSOFT v 8.0.0 (34,132); and 3) a pairwise comparison of PCs. This analysis led to the optimal K of 6. SNPs significantly correlated to these 6 PCs were identified through the Mahalanobis distance method as implemented in PCAdapt R (35) and the false discovery rate (FDR) of the p-values was calculated using the qvalue R package v2.18.0 (133). Outlier SNPs were extracted with the *get.pc* function of PCAdapt with an expected 5% of false positives (FDR, α=0.05), and their potential structural and/or functional effect was established implementing three tools: the SnpEff v4.3t program (131), the VariantAnnotation R package (135), and the ‘*annotate*’ function from bcftools using an *in-house* R script from a customized AaegL5 genome annotation file (see below).

To have an unbiased estimate of local adaptation, we established a statistical framework to associate the PC-specific outliers to a population. Briefly, we used the PC scores generated for each mosquito from the covariance matrix of the PCA, and then analyzed their distribution across populations on each of the 6 PCs, separately, as implemented in previous studies (136,137). A PC score represents not only the PCA projections of a single genome, but also an independent measure of variation in the form of values that are different from zero. We used these PC scores to identify members of a population with patterns of high variance to support their local genetic differentiation. We defined a population to be locally differentiated when the distribution of the PC scores significantly departs from zero. This significance was tested by evaluating whether PC scores from a population are significantly different from zero on the corresponding PC (one sample t-test, m=0), and also if they are significantly different from other populations (pairwise t-test). Final candidates were obtained after manually comparing those populations with significant PC scores. To support estimates of local genetic adaptation, we used two additional parameters of genetic diversity: F_ST_ and Tajima’s D, which were calculated using VCFTools. The F_ST_ value of each outlier SNP was used as a measurement of the “intensity” of genetic differentiation and Tajima’s D was estimated in genes with outlier SNPs to identify whether such genetic variation is different from neutrality (−0.5 < D <1.0).

To identify a set of genes that shows a strong signal of genetic differentiation between Aaa and Aaf, we first identified with a custom PERL program all genes associated with outlier SNPs and quantified their occurrence across genomic regions and effects (*i.e.*, synonymous and non-synonymous mutations). Next, allele frequencies of all non-synonymous SNPs were obtained for each gene across all populations. To evaluate significant differences between Aaa and Aaf populations in this filtered set of genes, we created 4 groups of populations based on the genetic diversity and divergence patterns we observed in our data: 1) THI and NGY, 2) RABs and RABd, 3) the rest of African populations, and 4) all out of Africa populations. Despite the genetic difference between these groups, our null hypothesis is to find genes significantly differentiated between Aaa and Aaf (Ho), this will be rejected (p>0.05) for cases in which no major differences can be detected using an ANOVA test.

A GO enrichment analysis across the three major GO categories (Biological Processes, Molecular Functions, and Cellular Components) was performed to identify functional groups that were enriched in this set of genes. Briefly, we used a custom genome database for *Ae. aegypti* with GO annotations (*in-house* org.Aaegypti.eg.db R package, see below) and the clusterProfiler R package v4.2.2 (138) to calculate the GO enrichment. *P*-values (p≤0.05) were corrected for multiple tests using the Benjamini-Hochberg procedure, and redundancy of enriched GO terms on each major GO classification was removed with the function *simplify*, both implemented in clusterProfiler (135). The custom org.Aaegypti.eg.db R package was built based on a collection of GO annotations retrieved from VectorBase version 59 (139) and a bioinformatic approach using BLAST (NCBI Diptera *non-redundant* (nr) database v5) and InterProScan v5 (140) scanning different protein domain databases: Pfam v33.1 (141), ProSiteProfiles v20.2 (142), SUPERFMILY v2.0 (143), and TIGRFAM v15.0 (144). Outlier SNPs were also mapped against a target set of 1132 genes, including 198 detoxification genes, 198 chemosensory genes (OR, IR and GR receptors), 391 immunity genes, 292 proteases, and 53 genes associated to multiple functions (32,64,65).

### Assessing Ka/Ks ratio

We estimated the ratio of non-synonymous (Ka) to synonymous (Ks) substitutions (also known as dN/dS ratio) across 13,503 of the 14,677 *Ae. aegypti* protein coding genes, and identified genes as evolving neutrally or nearly neutral with a conservative threshold of 0.95≥Ka/Ks≤1.05, or under negative selection when Ka/Ks<0.95 (and 0.80≥Ka/Ks<0.95 for weak negative selection), or under positive selection when Ka/Ks>1.05 (and 1.05>Ka/Ks≤1.2 for weak positive selection) (57). A total of 1,174 genes were not analyzed due to either the absence of SNPs and/or the presence of multiples indels affecting the positions of codons. To do this, we first extracted codons associated to SNPs and classified into non-degenerate (L0) sites, 2-fold (L2) degenerated sites, and 4-fold (L4) degenerated sites. Next, transition (Ts) and transversion (Tv) changes were identified for each codon by comparing the alternative and reference alleles and obtaining the number of Ts at L0 (A0), L2 (A2), and L4 (A2) degenerated sites, as well as the number of Tv at L0 (B0), L2 (B2), and L4 (B2) degenerated sites. The rates of *Ka* and *Ks* site substitution were calculated using the improved Kimura-2 parameters (K2P) Li’s method (145,146).

We also estimated the ratio of Ka/Ks by gene in each population using the PAML package v4.10.6 (147). For this analysis, we derived 539,298 gene nucleotide alignments with the *vcf2fasta* package (https://github.com/santiagosnchez/vcf2fasta) from protein coding genes with at least 1 SNP in each population. Then, each alignment was translated into amino acids sequences with *transeq* from the EMBOSS package v6.6.0.0 (148), and their corresponding codon alignments were created with the *pal2nal.pl* program v14 (149). A total of 162 protein coding genes were removed from this analysis due to the presence of multiples indels affecting the positions of codons, resulting in a final dataset of 539,136 codon alignments. A ML phylogenetic tree was reconstructed for each codon alignment with *FastTree* v2.1 (150) and the GTR+GAMMA model. The *one-ratio model* (M0) was used to calculate the ka/ks ratio average for the whole gene over all branches in the phylogeny with the *codeml* program from the PAML package (147). The Ka/Ks ratio of a gene was the average Ka/Ks ratios calculated within the gene. PAML detected 2.5-fold times more sites under selection than the improved K2P Li’s method. However, ∼37% of the Ka/Ks ratios detected with PAML exhibit standard deviations >10, suggesting either a high divergence among individuals within a population or an overestimation of the Ka/Ks ratio *per* gene. The latter is more probable because the PAML-based Ka/Ks ratio method ignores SNPs segregating within a population and transient SNPs within divergent populations (151–153). Based on these results, we further analyzed and discussed only the Ka/Ks ratios estimated with the improved K2P Li’s method.

### Analysis of *Ae. aegypti* nrEVEs

The frequency of each of the 252 nrEVEs annotated in AaegL5 was established in each of the analyzed populations using the SVD pipeline (46,48,154). The same procedure was used to verify the occurrence of *Ae. aegypti* nrEVEs in 4 WGS dataset from *Ae. mascarensis* (10).

The Vy-PER (155 and ViR (48) pipelines were used to search for novel viral integrations using a viral database assembled in October 2020, which encompasses a total of 1778 taxids and 3677 nucleotide sequences of both DNA and RNA viruses. To build the viral database, viral taxids were extracted from NCBI virus (https://www.ncbi.nlm.nih.gov/labs/virus/vssi/#/) according to three different criteria: (i) main known arboviral genera; (ii) ISVs identified from a search of NCBI PubMed publications between 2015 and 2020 using the keyword ‘insect specific viruses’; (iii) viruses having Diptera as host. Nucleotide sequences for the accessions corresponding to the selected taxids were retrieved using the NCBI E-utility tool. After removing duplicates, sequences were clustered at 97% identity using CD-HIT (156). Sequences were BLAST searched against a database of conserved eukaryotic and bacterial ribosomal sequences developed for SortmeRNA (157) and of Diptera ribosomal sequences extracted from (158). Entire sequences or parts of sequences matching ribosomal DNA were removed or masked, respectively. Homopolymers and repeats were identified using a custom script and masked with Ns. The presence of novel nrEVEs was further tested in all WGS data used for SNP discovery using the ViR_LTFinder script (48) to confirm their widespread distribution in African vs out-of-Africa samples. A subset of 26 out of the 64 novel nrEVEs were molecularly validated by PCR and Sanger sequencing in the mosquito DNA samples that had been used for WGS using primers designed on the bioinformatic predictions generated by ViR. PCRs were performed in 50 mL of volume containing 25 mL of 2X DreamTaq Green PCR Master Mix (ThermoFisher), 2.5 mL of 10 mM forward and reverse primers, 1 mL of DNA diluted 1:10 and sterile water to volume. Reactions without template served as negative controls. PCR products were visualized through electrophoresis on 2% agarose gels stained with ethidium bromide. Since multiple bands were often observed, bands of expected size were cut from the gels and purified using the GeneJET PCR Purification Kit (Thermo Scientific), following the manufacturer’s protocol. Purified PCR products were sent for Sanger sequencing to Macrogen Europe (Netherlands) to confirm nrEVE sequence.

This version of the Manuscript has not supplementary material.

## Declarations

### Ethics approval and consent to participate

Not applicable

### Consent for publication

Not applicable

### Data and materials availability

main text or the supplementary materials include all data and information on data accessibility.

### Author contributions

conceptualization and resources: A. Lozada-Chavez and M. Bonizzoni; whole-genome sequencing: J.A. Souza-Neto, B.C. Carlos and M. Bonizzoni; SNPs identification, population structure and genome-wide codon selection analyses: A. Lozada-Chávez and I. Lozada-Chávez; identification and analyses of nrEVEs: U. Palatini, N. Alfano and M. Bonizzoni; molecular biology data: D. Sogliani and N. Alfano; samples acquisition: T. Degefa, M.V. Sharakhova, A. Badolo, J. Prachumsri, M. Casas-Martinez, B.C. Carlos, L. Lambrechts, J.A. Souza-Neto; figures preparation: A. Lozada-Chávez; N. Alfano; D. Sogliani, U. Palatini; manuscript writing team: A. Lozada-Chávez, I. Lozada-Chávez, S. Elfekih, T. Degefa, M.V. Sharakhova, A. Badolo, J. Prachumsri, M. Casas-Martinez, B. C. Carlos, L. Lambrechts, J.A. Souza-Neto and M. Bonizzoni.

## Acknowledgements

We would like to thank members of the Bonizzoni’s lab. for fruitful discussion. We thank the members of the Department of Zoonosis and Vector Control from the Municipality of Bebedouro (Brazil) for assistance with mosquito collections.

